## Supplementary data for "Transient growth factor expression via mRNA in lipid nanoparticles promotes hepatocyte cell therapy to treat murine liver diseases"

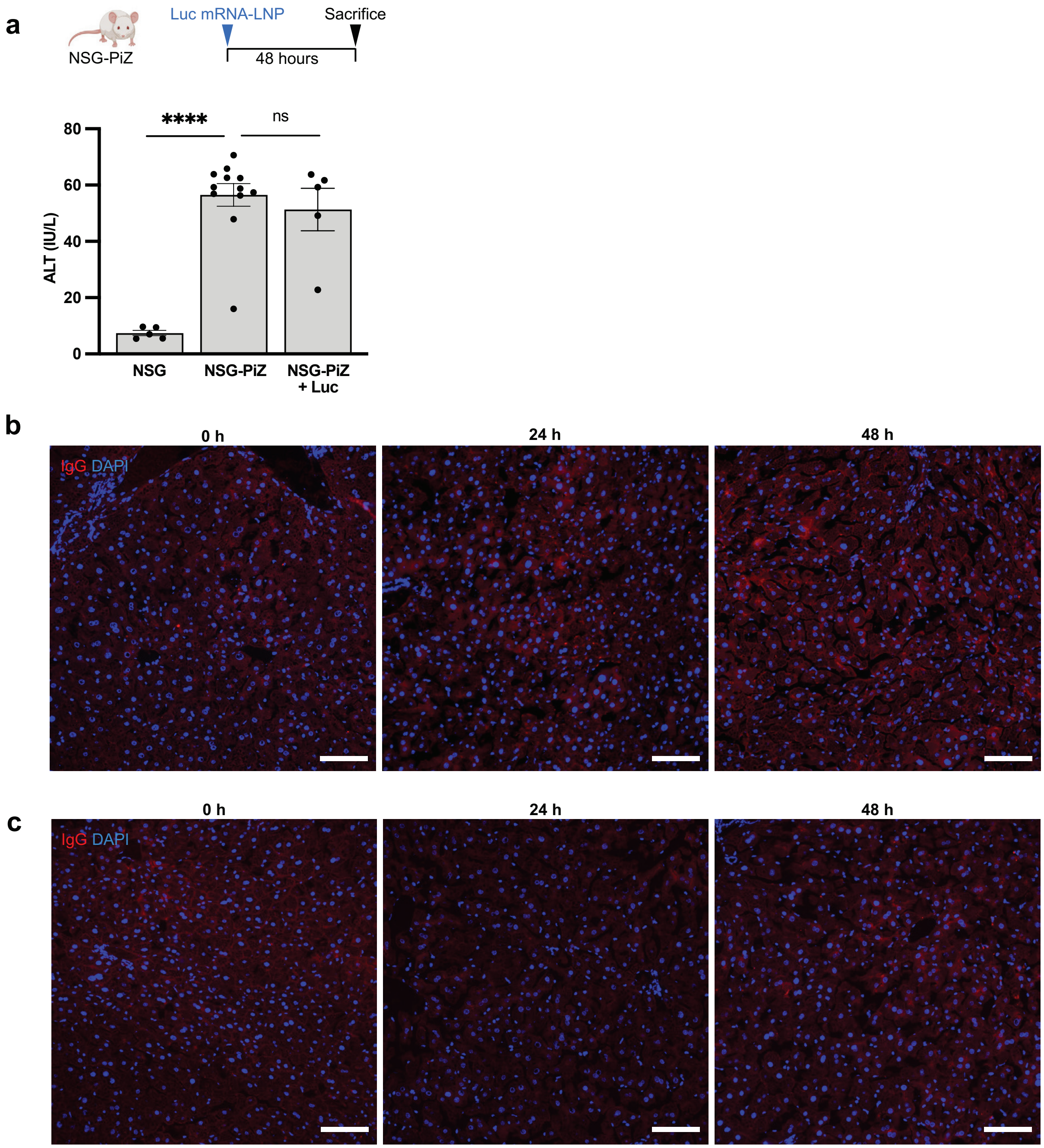


**Supplementary Fig. 1 | mRNA-LNP do not induce hepatotoxicity & isotype control staining for HGF and EGF antibodies. a** Liver enzyme alanine aminotransferase (ALT) levels in serum 48 hours post Luc mRNA-LNP injection measured via kinetic spectrophotometric assay. N=5 or 12. Each dot represents one mouse, error bars = SEM, ns $P > 0.05$, * $P \le 0.05$, ** $P \le 0.01$, *** $P \le 0.001$, **** $P \le 0.0001$. **b** Representative images of immunofluorescence staining for the HGF antibody isotype control on liver tissue of untreated NSG-PiZ mice and 24 hours or 48 hours after IV administration of HGF+EGF mRNA-LNP. **c** Representative images of immunofluorescence staining for the EGF antibody isotype control on liver tissue of untreated NSG-PiZ mice and 24 hours or 48 hours after IV administration of HGF+EGF mRNA-LNP. For panels b,c: N=3, scale = 100um.

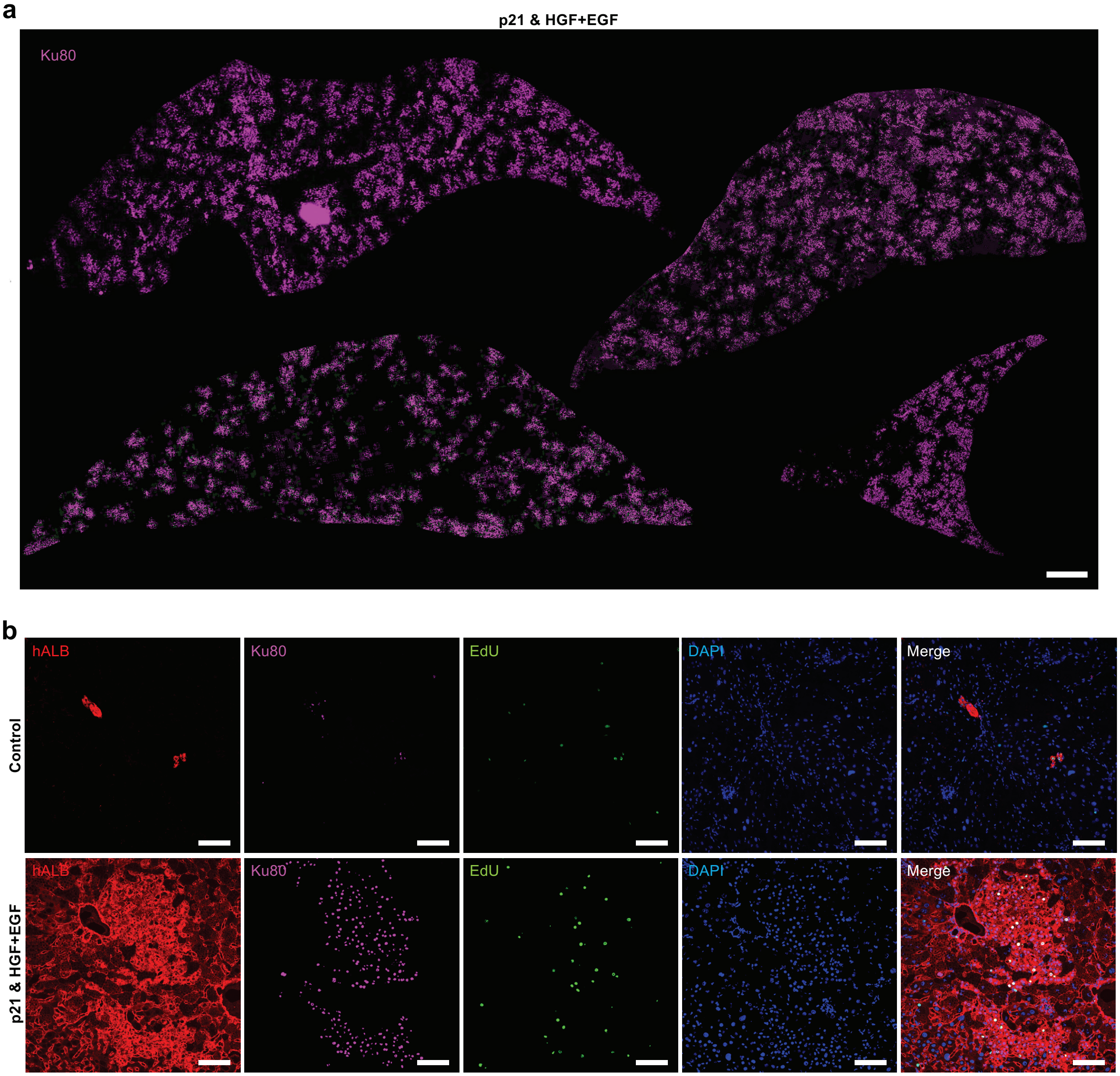


**Supplementary Fig. 2 | HGF+EGF mRNA-LNP treatments lead to sustained and robust engraftment of PHHs in p21/NSG-PiZ mice. a** Representative images of Ku80 immunofluorescence stain on liver sections from the experimental group 5 weeks post transplantation. One representative lobe is shown from each sample, various liver lobes are highlighted. N = 5, scale = 1000 um. **b** Representative images of hALB/Ku80/EdU immunofluorescence stain on liver sections from experimental group 5 weeks post transplantation. Split and merged channels are shown. N = 5, scale = 100um.

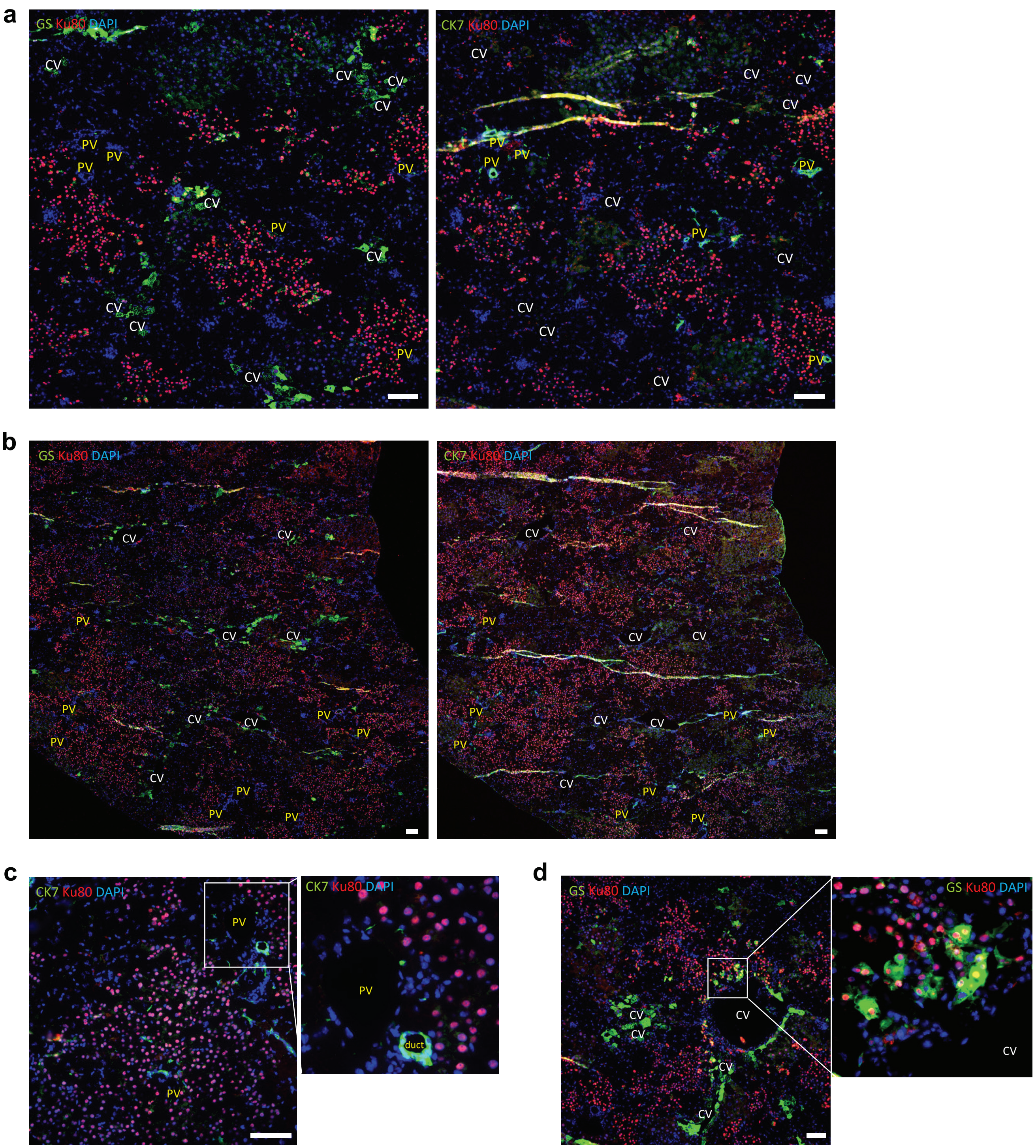


**Supplementary Fig. 3 | Engrafted PHHs originate in the periportal region, span all liver zones, and adopt appropriate zonation markers. a** Representative images of GS/Ku80 and CK7/Ku80 immunofluorescence stain on serial liver sections from experimental group 5 weeks post transplantation, highlighting an area of smaller clusters of engrafted PHHs. **b** Representative images of GS/Ku80 and CK7/Ku80 immunofluorescence stain on serial liver sections from experimental group 5 weeks post transplantation, highlighting an area of larger clusters of engrafted PHHs. **c** Representative image of CK7/Ku80 immunofluorescence stain on liver section from experimental group 5 weeks post transplantation, highlighting PHHs engrafted in periportal region. **d** Representative image of GS/Ku80 immunofluorescence stain on liver section from experimental group 5 weeks post transplantation, highlighting $GS^{+}Ku80^{+}$ PHHs engrafted in pericentral region. For all panels: Portal vein (PV) regions are annotated in yellow, and central vein (CV) regions are annotated in white. N = 5, scale = 100um.

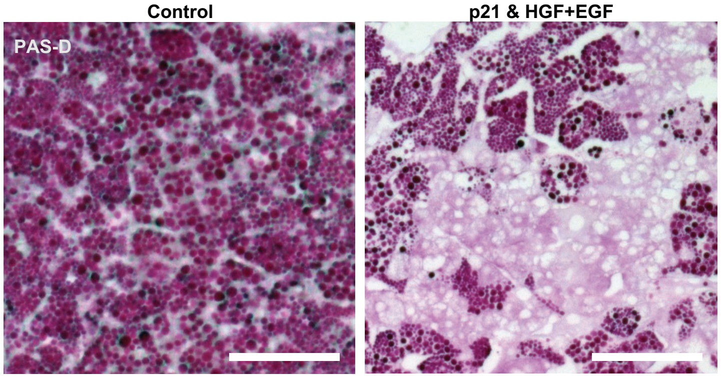


**Supplementary Fig. 4 | PHH engraftment levels in experimental group are sufficient to reduce Z-AAT burden in the liver.** Representative images of periodic acid Schiff - diastase (PAS-D) histology stain on liver sections from control and experimental groups 5 weeks post transplantation. hZ-AAT polymers are not digested by diastase, stain PAS-D$^+$, and are dark magenta in color. For all panels: N = 5, scale = 100um.

**Supplementary Table 1:** DNA sequences to generate nucleoside-modified mRNAs

| Name | Sequence |
|---|---|
| Luciferase | ATGGAGGACGCCAAGAACATCAAGAAGGGCCCCGCCCCCTTCTACCCCCTGGA<br>GGACGGCACCGCCGGCGAGCAGCTGCACAAGGCCATGAAGCGgTACGCCCTG<br>GTGCCCGGCACCATCGCCTTCACCGACGCCCACATCGAGGTGGACATCACCTA<br>CGCCGAGTACTTCGAGATGTCCGTGCGCCTGGCCGAGGCCATGAAGCGgTACG<br>GCCTGAACACCAACCACCGCATCGTGGTGTGCTCCGAGAACTCCCTGCAGTTC<br>TTCATGCCCGTGCTGGGCGCCCTGTTCATCGGCGTGGCCGTGGCCCCCGCCA<br>ACGACATCTACAACGAGCGCGAGCTGCTGAACTCCATGGGCATCTCCCAGCCC<br>ACCGTGGTGTTCGTGTCCAAGAAGGGCCTGCAGAAGATCCTGAACGTGCAGAA<br>GAAGCTGCCCATCATCCAGAAGATCATCATCATGGACTCCAAGACCGACTACCA<br>GGGCTTCCAGTCCATGTACACCTTCGTGACCTCCCACCTGCCCCCCGGCTTCA<br>ACGAGTACGACTTCGTGCCCGAGTCCTTCGACCGCGACAAGACCATCGCCCTG<br>ATCATGAACTCCTCCGGCTCCACCGGCCTGCCCAAGGGCGTGGCCCTGCCCCA<br>CCGCACCGCCTGCGTGCGCTTCTCCCACGCCCGCGACCCCATCTTCGGCAACC<br>AGATCATCCCCGACACCGCCATCCTGTCCGTGGTGCCCTTCCACCACGGCTTC<br>GGCATGTTCACCACCCTGGGCTACCTGATCTGCGGCTTCCGCGTGGTGCTGAT<br>GTACCGCTTCGAGGAGGAGCTGTTCCTGCGCTCCCTGCAGGACTACAAGATCC<br>AGTCCGCCCTGCTGGTGCCCACCCTGTTCTCCTTCTTCGCCAAGTCCACCCTG<br>ATCGACAAGTACGACCTGTCCAACCTGCACGAGATCGCCTCCGGCGGCGCCCC<br>CCTGTCCAAGGAGGTGGGCGAGGCCGTGGCCAAGCGgTTCCACCTGCCCGGC<br>ATCCGCCAGGGCTACGGCCTGACCGAGACCACCTCCGCCATCCTGATCACCCC<br>CGAGGGCGACGACAAGCCCGGCGCCGTGGGCAAGGTGGTGCCCTTCTTCGAG<br>GCCAAGGTGGTGGACCTGGACACCGGCAAGACCCTGGGCGTGAACCAGCGCG<br>GCGAGCTGTGCGTGCGCGGCCCCATGATCATGTCCGGCTACGTGAACAACCCC<br>GAGGCCACCAACGCCCTGATCGACAAGGACGGCTGGCTGCACTCCGGCGACA<br>TCGCCTACTGGGACGAGGACGAGCACTTCTTCATCGTGGACCGCCTGAAGTCC<br>CTGATCAAGTACAAGGGCTACCAGGTGGCCCCCGCCGAGCTGGAGTCCATCCT<br>GCTGCAGCACCCCAACATCTTCGACGCCGGCGTGGCCGGCCTGCCCGACGAC<br>GACGCCGGCGAGCTGCCCGCCGCCGTGGTGGTGCTGGAGCACGGCAAGACC<br>ATGACCGAGAAGGAGATCGTGGACTACGTGGCCTCCCAGGTGACCACCGCCAA<br>GAAGCTGCGCGGCGGCGTGGTGTTCGTGGACGAGGTGCCCAAGGGCCTGACC<br>GGCAAGCTGGACGCCCGCAAGATCCGCGAGATCCTGATCAAGGCCAAGAAGG<br>GCGGCAAGATCGCCGTG |
| eGFP | ATGGTGAGCAAGGGCGAGGAGCTGTTCACCGGGGTGGTGCCCATCCTGGTCG<br>AGCTGGACGGCGACGTAAACGGCCACAAGTTCAGCGTGTCCGGCGAGGGCGA<br>GGGCGATGCCACCTACGGCAAGCTGACCCTGAAGTTCATCTGCACCACCGGCA<br>AGCTGCCCGTGCCCTGGCCCACCCTCGTGACCACCCTGACCTACGGCGTGCA<br>GTGCTTCAGCCGCTACCCCGACCACATGAAGCAGCACGACTTCTTCAAGTCCG<br>CCATGCCCGAAGGCTACGTCCAGGAGCGCACCATCTTCTTCAAGGACGACGGC<br>AACTACAAGACCCGCGCCGAGGTGAAGTTCGAGGGCGACACCCTGGTGAACC<br>GCATCGAGCTGAAGGGCATCGACTTCAAGGAGGACGGCAACATCCTGGGGCAC<br>AAGCTGGAGTACAACTACAACAGCCACAACGTCTATATCATGGCCGACAAGCAG<br>AAGAACGGCATCAAGGTGAACTTCAAGATCCGCCACAACATCGAGGACGGCAG<br>CGTGCAGCTCGCCGACCACTACCAGCAGAACACCCCCATCGGCGACGGCCCC<br>GTGCTGCTGCCCGACAACCACTACCTGAGCACCCAGTCCGCCCTGAGCAAAGA<br>CCCCAACGAGAAGCGCGATCACATGGTCCTGCTGGAGTTCGTGACCGCCGCCG<br>GGATCACTCTCGGCATGGACG AGCTGTACAAG |

| | |
|---|---|
| HGF | ATGTGGGTGACCAAGCTGCTGCCCGCCCTGCTGCTGCAGCACGTGCTGCTGCA<br>CCTGCTGCTGCTGCCCATCGCCATCCCCTACGCCGAGGGCCAGCGCAAGCGC<br>CGCAACACCATCCACGAGTTCAAGAAGTCCGCCAAGACCACCCTGATCAAGATC<br>GACCCCGCCCTGAAGATCAAGACCAAGAAGGTGAACACCGCCGACCAGTGCG<br>CCAACCGCTGCACCCGCAACAAGGGCCTGCCCTTCACCTGCAAGGCCTTCGTG<br>TTCGACAAGGCCCGCAAGCAGTGCCTGTGGTTCCCCTTCAACTCCATGTCCTCC<br>GGCGTGAAGAAGGAGTTCGGCCACGAGTTCGACCTGTACGAGAACAAGGACTA<br>CATCCGCAACTGCATCATCGGCAAGGGCCGCTCCTACAAGGGCACCGTGTCCA<br>TCACCAAGTCCGGCATCAAGTGCCAGCCCTGGTCCTCCATGATCCCCCACGAG<br>CACTCCTTCCTGCCCTCCTCCTACCGCGGCAAGGACCTGCAGGAGAACTACTG<br>CCGCAACCCCCGCGGCGAGGAGGGCGGCCCCTGGTGCTTCACCTCCAACCCC<br>GAGGTGCGCTACGAGGTGTGCGACATCCCCCAGTGCTCCGAGGTGGAGTGCAT<br>GACCTGCAACGGCGAGTCCTACCGCGGCCTGATGGACCACACCGAGTCCGGC<br>AAGATCTGCCAGCGgTGGGACCACCAGACCCCCCACCGCCACAAGTTCCTGCC<br>CGAGCGgTACCCCGACAAGGGCTTCGACGACAACTACTGCCGCAACCCCGACG<br>GCCAGCCCCGCCCCTGGTGCTACACCCTGGACCCCCACACCCGCTGGGAGTA<br>CTGCGCCATCAAGACCTGCGCCGACAACACCATGAACGACACCGACGTGCCCC<br>TGGAGACCACCGAGTGCATCCAGGGCCAGGGCGAGGGCTACCGCGGCACCGT<br>GAACACCATCTGGAACGGCATCCCCTGCCAGCGgTGGGACTCCCAGTACCCCC<br>ACGAGCACGACATGACCCCCGAGAACTTCAAGTGCAAGGACCTGCGCGAGAAC<br>TACTGCCGCAACCCCGACGGCTCCGAGTCCCCCTGGTGCTTCACCACCGACCC<br>CAACATCCGCGTGGGCTACTGCTCCCAGATCCCCAACTGCGACATGTCCCACG<br>GCCAGGACTGCTACCGCGGCAACGGCAAGAACTACATGGGCAACCTGTCCCAG<br>ACCCGCTCCGGCCTGACCTGCTCCATGTGGGACAAGAACATGGAGGACCTGCA<br>CCGCCACATCTTCTGGGAGCCCGACGCCTCCAAGCTGAACGAGAACTACTGCC<br>GCAACCCCGACGACGACGCCCACGGCCCCTGGTGCTACACCGGCAACCCCCT<br>GATCCCCTGGGACTACTGCCCCATCTCCCGCTGCGAGGGCGACACCACCCCCA<br>CCATCGTGAACCTGGACCACCCCGTGATCTCCTGCGCCAAGACCAAGCAGCTG<br>CGCGTGGTGAACGGCATCCCCACCCGCACCAACATCGGCTGGATGGTGTCCCT<br>GCGCTACCGCAACAAGCACATCTGCGGCGGCTCCCTGATCAAGGAGTCCTGGG<br>TGCTGACCGCCCGCCAGTGCTTCCCCTCCCGCGACCTGAAGGACTACGAGGC<br>CTGGCTGGGCATCCACGACGTGCACGGCCGCGGCGACGAGAAGTGCAAGCAG<br>GTGCTGAACGTGTCCCAGCTGGTGTACGGCCCCGAGGGCTCCGACCTGGTGC<br>TGATGAAGCTGGCCCGCCCCGCCGTGCTGGACGACTTCGTGTCCACCATCGAC<br>CTGCCCAACTACGGCTGCACCATCCCCGAGAAGACCTCCTGCTCCGTGTACGG<br>CTGGGGCTACACCGGCCTGATCAACTACGACGGCCTGCTGCGCGTGGCCCACC<br>TGTACATCATGGGCAACGAGAAGTGCTCCCAGCACCACCGCGGCAAGGTGACC<br>CTGAACGAGTCCGAGATCTGCGCCGGCGCCGAGAAGATCGGCTCCGGCCCCT<br>GCGAGGGCGACTACGGCGGCCCCCTGGTGTGCGAGCAGCACAAGATGCGCAT<br>GGTGCTGGGCGTGATCGTGCCCGGCCGCGGCTGCGCCATCCCCAACCGCCCC<br>GGCATCTTCGTGCGCGTGGCCTACTACGCCAAGTGGATCCACAAGATCATCCTG<br>ACCTACAAGGTGCCCCAGTCCTAA |
| EGF (secreted) | ATGGCCACCGGCTCCCGCACCTCCCTGCTGCTGGCCTTCGGCCTGCTGTGCCT<br>GCCCTGGCTGCAGGAGGGCTCCGCCATGAACTCCGACTCCGAGTGCCCCCTG<br>TCCCACGACGGCTACTGCCTGCACGACGGCGTGTGCATGTACATCGAGGCCCT<br>GGACAAGTACGCCTGCAACTGCGTGGTGGGCTACATCGGCGAGCGgTGCCAGT<br>ACCGCGACCTGAAGTGGTGGGAGCTGCGCTAA |

**Supplementary Table 2:** List of antibodies

| Antibody | Company | Catalog # | Species | Application | Dilution | Notes |
|---|---|---|---|---|---|---|
| GFP | Invitrogen | A10262 | Chicken | IF-Fr | 1:300 | - |
| HGF | R&D | AF-294 | Goat | IF-Fr | 1:20 | - |
| EGF | R&D | MAB236 | Mouse | IF-Fr | 1:30 | M.O.M. Kit (Vector Labs BMK-2202) |
| P21 | Abcam | ab188224 | Rabbit | IF-Fr | 1:500 | 0.2% triton X with primary antibody incubation |
| hALB | Bethyl | A80-129A | Goat | IF-Fr | 1:1000 | - |
| Ku80 | Cell Signaling | mAb #2180 | Rabbit | IF-Fr | 1:50 | 0.2% triton X with primary antibody incubation |
| 2C1 | Hycult Biotech | HM2289 | Mouse | IF-Fr | 1:50 | M.O.M. Kit (Vector Labs BMK-2202) |

IF-Fr: immunofluorescence, frozen sections
